# Response Adjusted for Days of Antibiotic Risk (RADAR): evaluation of a novel method to analyze antibiotic stewardship interventions

**DOI:** 10.1101/107300

**Authors:** Valentijn A. Schweitzer, Maarten van Smeden, Douwe F. Postma, Jan Jelrik Oosterheert, Marc J.M. Bonten, Cornelis H. van Werkhoven

**Affiliations:** Julius Center for Health Sciences and Primary Care, University Medical Centre Utrecht, Utrecht, The Netherlands; Departments of Internal Medicine and Infectious Diseases, University Medical Centre Utrecht, Utrecht, The Netherlands; Department of Medical Microbiology, University Medical Centre Utrecht, Utrecht, The Netherlands

**Author notes:** Corresponding author: Valentijn A. Schweitzer, MD, Julius Center for Health Sciences and Primary Care, University Medical Center Utrecht, Heidelberglaan 100, Post Box 85500, Utrecht 3508 GA, The Netherlands. Alternate corresponding author: Cornelis H. van Werkhoven, MD, Julius Center for Health Sciences and Primary Care, University Medical Center Utrecht, Heidelberglaan 100, Post Box 85500, Utrecht 3508 GA, The Netherlands.

**Keywords:** antibiotic stewardship, methodology, DOOR/RADAR, antibiotic use strategies, community-acquired pneumonia

## Abstract

**OBJECTIVES:** The Response Adjusted for Days of Antibiotic Risk (RADAR)-statistic was proposed to improve efficiency of antibiotic stewardship trials. We studied the behavior of RADAR in a non-inferiority trial in which a beta-lactam monotherapy strategy (BL, n=656) was non-inferior to fluoroquinolone monotherapy (FQL, n=888) for moderately-severe community-acquired pneumonia (CAP) patients.

**METHODS:** Patients were ranked according to clinical outcome, using five or eight categories, and antibiotic use. RADAR was calculated as the probability that the BL group had a more favorable ranking than the FQL group. To investigate the sensitivity of RADAR to detrimental clinical outcome we simulated increasing rates of 90-day mortality in the BL group and performed the RADAR and non-inferiority analysis.

**RESULTS:** The RADAR of the BL-group compared to the FQL group was 60.3% (95% confidence interval 57.9%-62.7%) using five and 58.4% (95% CI 56.0%-60.9%) using eight clinical outcome categories, all in favor of BL. Sample sizes for RADAR were 250 and 580 patients per study arm using five or eight clinical outcome categories, respectively, reflecting 38% and 89% of the original non-inferiority sample size calculation. With simulated mortality rates, loss of non-inferiority of the BL-group occurred at a relative risk of 1.125 in the conventional analysis, whereas using RADAR the BL-group lost superiority at a relative risk of mortality of 1.25 and 1.5, with eight and five clinical outcome categories, respectively.

**CONCLUSIONS:** RADAR favored BL over FQL therapy for CAP. Although RADAR required fewer patients than conventional non-inferiority analysis, the statistic was less sensitive to detrimental outcomes.

## INTRODUCTION

Antibiotic resistance is associated with prolonged hospital stay and increased mortality and healthcare costs^1,2^. Selection of antibiotic resistant bacteria is facilitated through use of antibiotics^3^. Previous studies found high rates of inappropriate antibiotic use in different clinical settings^4–6^. With antibiotic stewardship programs, physicians aim to ensure therapeutic efficacy while limiting adverse events of antibiotic overuse, such as the emergence of resistance, adverse drug events and costs^3,4^. In general, trials to evaluate antibiotic stewardship interventions are designed to show an increase in appropriate antibiotic use without compromising clinical outcome, frequently using a non-inferiority design. However, non-inferiority trials may suffer from analysis of suboptimal or subjective clinical outcomes, such as “clinical cure”, and are frequently underpowered to demonstrate non-inferiority for objective clinical outcomes such as mortality^7^. Also, comparing benefit and harm can be difficult, for example when an intervention decreases antibiotic use at the cost of more complications. Finally, non-inferiority trials usually require large numbers of subjects and choosing the optimal non-inferiority margin may be subjective and may lead to discussion after completion of the trial^8,9^.

In search for a better method to analyze antibiotic stewardship interventions, the Response Adjusted for Days of Antibiotic Risk (RADAR) statistic was proposed^10^. For its computation, patients are first classified based on mutually exclusive, hierarchical levels corresponding to clinical outcome of the patient, e.g. complication free survival, survival with complications and mortality. Within each clinical outcome category, patients are subcategorized according to their level of antibiotic use. All patients are then ranked according to their category, where patients with a better clinical outcome, receiving less antibiotics or antibiotics with a narrower antimicrobial spectrum have a more favorable rank. Theoretically, through this ordinal classification, patients with a worse clinical outcome always have a lower ranking, while within outcome categories, appropriateness of antibiotic consumption determines the ranking. In the analysis, the distributions of the rankings pre and post antibiotic stewardship intervention are compared by combining clinical outcomes and antibiotic use, RADAR allows to analyze antibiotic stewardship interventions as superiority instead of non-inferiority trials, thereby requiring a smaller sample size^10,11^. However, RADAR also may have disadvantages, for example: choosing the components of the hierarchical clinical categories is subjective, RADAR results in a percentage which is difficult to interpret and it is uncertain whether clinical safety can be demonstrated with the reduced sample size^12,13^. Therefore, RADAR needs to be evaluated with real-life clinical trial data. For this purpose, we used data from a non-inferiority trial of empirical antibiotic treatment strategies in patients with community-acquired pneumonia (CAP) admitted to non-intensive care unit (non-ICU) wards^14^. We study the application of RADAR in comparison to the conventional non-inferiority analysis to determine the sensitivity to choices in the analysis by (1) calculating RADAR with different clinical outcome categories and different levels of antibiotic use, (2) quantifying the effect of sample size on the certainty of the outcome with both methods, and (3) determining the sensitivity of RADAR and non-inferiority to worse clinical outcomes by simulations.

## METHODS

### Data collection

The Community-Acquired Pneumonia — Study on the Initial Treatment with Antibiotics of Lower Respiratory Tract Infections (CAP-START) was performed between February 2011 and August 2013 in 7 hospitals in the Netherlands^14,15^. In the CAP-START trial, patients above 18 years of age who were admitted to a non-ICU ward for suspicion of CAP were eligible for study participation. Hospitals participating in the trial were cluster-randomized to beta-lactam monotherapy (BL), beta-lactam with a macrolide (BLM), or fluoroquinolone monotherapy (FQL) as preferred empiric treatment strategies for a 4 months period in random order. Physicians in the participating hospitals were repeatedly reminded of the current antibiotic strategy by local investigators with the use of newsletters and presentations to ensure strategy adherence. Deviation from the assigned treatment or subsequent change was allowed when medically indicated. Depending on the study arm, adherence to the strategy varied between 70 to 80%.

### Clinical outcome rankings

For the current analysis we compared the BL group to the FQL group, considering the latter as the control group with a high proportion of patients receiving antibiotics covering atypical pathogens. Implementing BL was considered the antibiotic stewardship intervention aiming to reduce the use of atypical coverage. Clinical outcome categories were constructed, in accordance with RADAR recommendations, with either two, five, or eight mutually exclusive hierarchical levels for clinical outcome (Table 1)^10^. RADAR is not intended to be used with a simple binary clinical outcome ranking^10^. We only use the 2-category RADAR to illustrate the behavior of RADAR in response to the amount of clinical categories used. Categories of antibiotic use were defined either as receipt of any atypical coverage (less favorable ranking) at any time during hospitalization versus no atypical coverage (more favorable ranking), or as the number of in-hospital days on atypical coverage (more days equals less favorable ranking). Atypical coverage was defined as antibiotic treatment with azithromycin, erythromycin, clarithromycin, moxifloxacin, levofloxacin, doxycyclin or ciprofloxacin. RADAR is a rank-order statistic which reflects the probability that a randomly selected patient assigned to the intervention group (here the BL group) has a more favorable ranking compared to a randomly selected patient from the control group (here the FQL group)^10^. A RADAR-statistic of 50% implies that the BL and FQL group are equally ranked, <50% indicates that either the clinical outcome and/or the antibiotic use is worse in the BL group and >50% indicates that either the clinical outcome and/or the antibiotic use is better in the BL group. Two thousand bootstrap samples were generated to estimate 95% confidence intervals for the RADAR-statistic^10,16^. Statistical significance was declared when the confidence interval did not overlap a predefined clinically relevant value (55% as proposed by Evans et al.). For simplicity of illustration, cluster effects arising from the cluster-randomized design of the study were ignored in the calculation of RADAR; the intra-cluster correlation for 90-day mortality was estimated to be 4.5E-7 and is therefore negligible^14^. All analyses were performed using R software, version 3.0.2 (R Project for Statistical Computing)^17^.

**Table 1:**
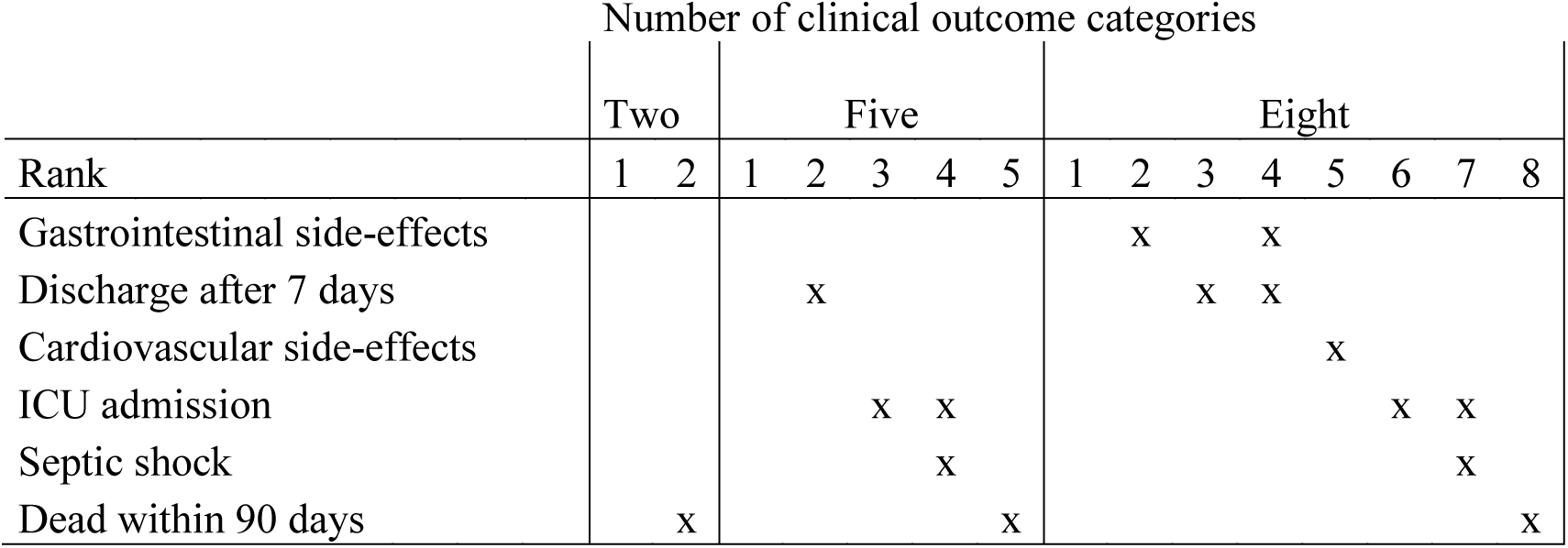
The constructed clinical outcome categories based on two, five and eight mutually exclusive hierarchical levels and corresponding ranks.

### Effect of sample size on non-inferiority analysis for mortality

The CAP-START trial was designed to demonstrate non-inferiority on day-90 mortality, with an predicted mortality rate of 5%, a non-inferiority margin of 3%, a one-sided alpha of 0.05, a power of 80%, and negligible intra-cluster correlation, yielding a required sample size of 650 per study arm^14,15^. The observed day-90 mortality rate was 10%. When re-estimating the sample size assuming 10% mortality, the required sample size is 1126 per study arm. The required sample size for RADAR was calculated to demonstrate a probability of at least 55% that a patient assigned to the BL group had a more favorable ranking than a patient assigned to the FQL group with a two-sided alpha of 0.05 and a power of 80%. The RADAR sample size was calculated using a Wilcoxon-Mann-Whitney test both with the predicted mortality of 5% and the observed mortality of 10%. For the sample size calculation using the observed mortality, the RADAR rank distributions were determined with the clinical outcomes and antibiotic use as observed in the CAP-START trial (Table 2). For the sample size calculation with the predicted mortality, we assumed that the distribution of remaining clinical outcomes and antibiotic use of the patients was the same as observed. Sample size calculations were confirmed with power simulations (results not shown).

**Table 2:** Distribution of rankings* in the BL period compared to the FQL period with either two or five clinical outcome categories.

|  |  | Two clinical<br>outcome categories |  | Five clinical outcome<br>categories |  | Eight clinical outcome<br>categories |  |
| --- | --- | --- | --- | --- | --- | --- | --- |
| Rank | Any<br>atypical<br>coverage | BL<br>N(%) | FQL<br>N(%) | BL<br>N(%) | FQL<br>N(%) | BL<br>N(%) | FQL<br>N (%) |
| 1 | - | 366 (56.0) | 80 (9.0) | 232 (35.5) | 48 (5.4) | 220 (33.6) | 44 (5.0) |
|  | + | 229 (35.0) | 729 (82.2) | 116 (17.7) | 445 (50.2) | 109 (16.7) | 423 (47.7) |
| 2 | - | 36 (5.5) | 12 (1.4) | 134 (20.5) | 32 (3.6) | 8 (1.2) | 3 (0.3) |
|  | + | 23 (3.5) | 66 (7.4) | 104 (15.9) | 275 (31.0) | 6 (0.9) | 13 (1.5) |
| 3 | - | - | - | 0 (0.0) | 0 (0.0) | 106 (16.2) | 26 (2.9) |
|  | + | - | - | 6 (0.9) | 8 (0.9) | 87 (13.3) | 234 (26.4) |
| 4 | - | - | - | 0 (0.0) | 0 (0.0) | 8 (1.2) | 4 (0.5) |
|  | + | - | - | 3 (0.5) | 1 (0.1) | 6 (0.9) | 8 (0.9) |
| 5 | - | - | - | 36 (5.5) | 12 (1.4) | 24 (3.7) | 3 (0.3) |
|  | + | - | - | 23 (3.5) | 66 (7.4) | 12 (1.8) | 27 (3.0) |
| 6 | - | - | - |  |  | 0 (0.0) | 0 (0.0) |
|  | + | - | - |  |  | 6 (0.9) | 8 (0.9) |
| 7 | - | - | - |  |  | 0 (0.0) | 0 (0.0) |
|  | + | - | - |  |  | 3 (0.5) | 1 (0.1) |
| 8 | - | - | - |  |  | 36 (5.5) | 12 (1.4) |
|  | + | - | - |  |  | 23 (3.5) | 66 (7.4) |

\* The definitions of the clinical outcome categories are shown in table 1.

To assess the impact of reduced RADAR sample size on non-inferiority of clinical outcome, we analyzed the risk difference of 90-day mortality if the amount of patients required for the RADAR analysis were included in the trial.

### Sensitivity of RADAR-statistic and the non-inferiority analysis for clinical outcome

The sensitivity of RADAR and the non-inferiority analysis for clinical outcome was tested by simulating increased mortality in the BL-group. In each simulation, a random selection of patients from the BL-group were reclassified to the “death within 90 days” clinical outcome category. The simulated RADAR for five and eight clinical categories, and the mortality risk differences between BL and FQL were plotted against the simulated relative risk of mortality. To determine the sensitivity of the statistics for clinical outcome, we used the simulated relative risk at which superiority (for RADAR) or non-inferiority (for the conventional analysis) was lost.

## RESULTS

### RADAR outcome of the CAP-START trial

From the 2,283 patients included in the CAP-START study, 656 were assigned to BL and 888 to FQL. When defining antibiotic use as ‘any atypical coverage’, the RADAR of the BL-group compared to the FQL group was 71.5% (95% confidence interval 69.2%-73.7%) using two clinical outcome categories, 60.3% (95% confidence interval 57.9%-62.7%) using five clinical outcome categories, and 58.4% (95% confidence interval 56.0%-60.9%) using eight clinical outcome categories, all in favor of BL (Figure 1). The RADAR-statistics were comparable when using ‘days of atypical coverage’ (Figure 1).

**Figure 1:**
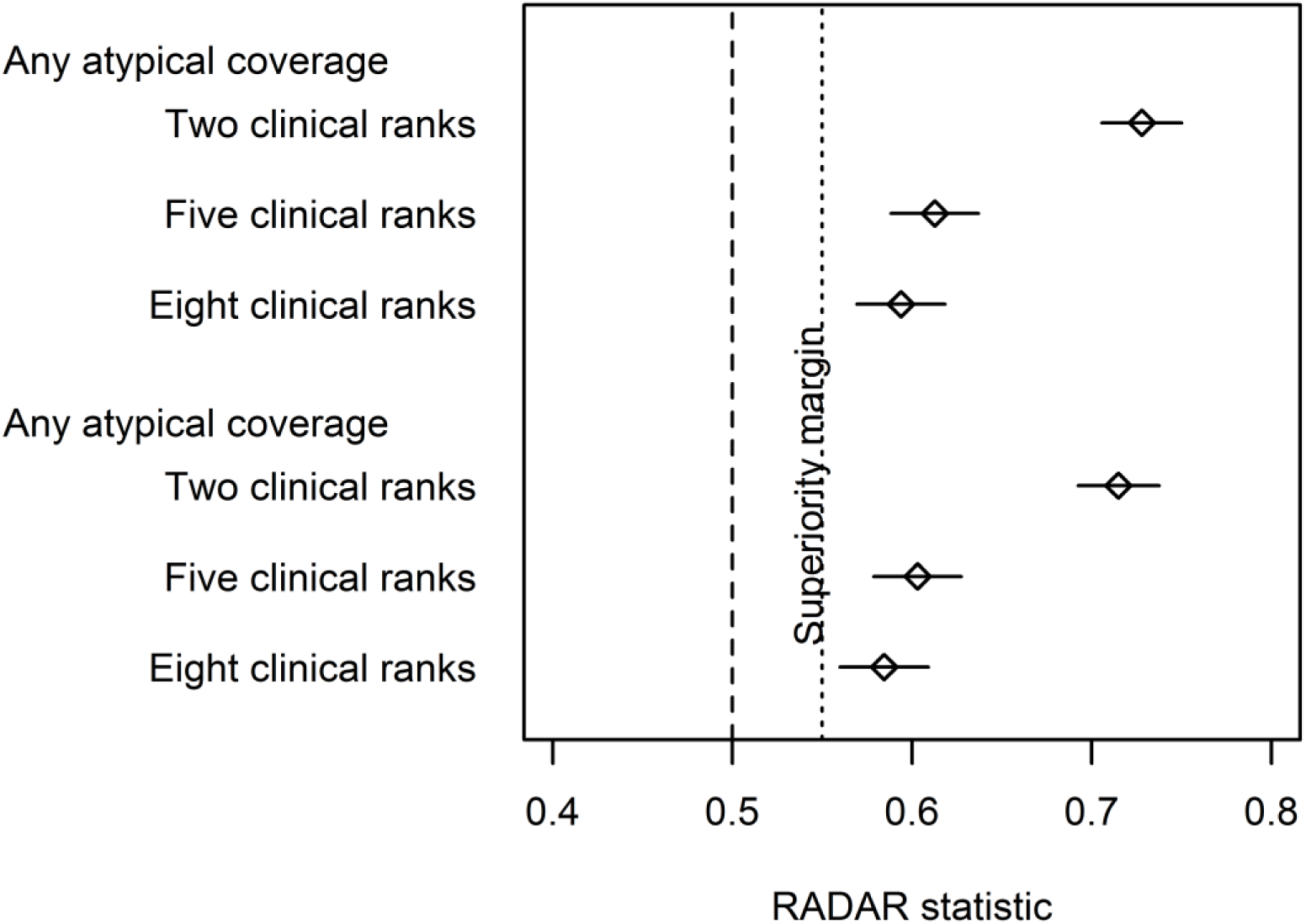
RADAR outcome of the CAP-START trial with two, five or eight clinical outcome categories and with antibiotic use defined as either any atypical coverage during hospitalization or days of atypical coverage received.

### Effect of sample size on non-inferiority analysis of mortality

The calculated required sample size for RADAR using five clinical categories to demonstrate superiority with the lower bound of the confidence interval above 55% was 250 patients per study arm, i.e. 38% of the non-inferiority sample size of the original CAP-START study using the predicted 5% mortality or 360 patients per study arm, i.e. 32% of the non-inferiority sample size using the observed 10% mortality (Table 3). The required sample size for the RADAR-analysis using eight clinical categories was 580 per study arm, i.e. 89% of the non-inferiority sample size using the predicted 5% mortality or 875 per study arm, i.e. 78% of the non-inferiority sample size using the observed 10% mortality. Based on the RADAR sample size calculation with five clinical outcome categories, 250 patients per study arm, or 500 consecutive patients enrolled which would yield a risk difference in 90-day mortality of −0.21% (-4.1% to 3.6%, 95% confidence interval). The required sample size with eight clinical outcome categories would correspond to 580 patients per study arm, which would result in a risk difference of −0.06% (-2.6% to 2.4%, 95% confidence interval). The corresponding power to demonstrate non-inferiority would be 45.7% when using five clinical outcome categories and 75.8% using eight clinical outcome categories. This demonstrates that, when using five clinical outcome categories, RADAR would allow a marked reductions in sample size but the method would have insufficient power to demonstrate non-inferiority on 90-day mortality. In contrast, when using eight clinical outcome categories, RADAR requires a similar sample size and consequently would have similar power to demonstrate non-inferiority on 90-day mortality.

**Table 3:**
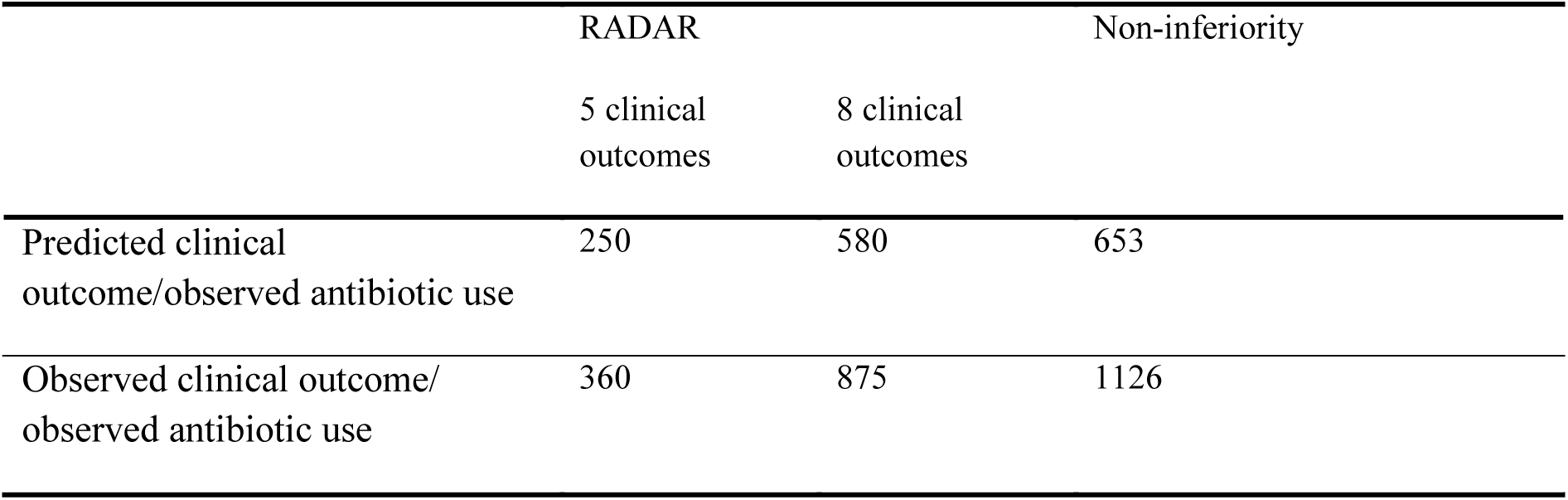
Estimated required sample size per study arm for RADAR and the non-inferiority design

### Sensitivity of RADAR-statistic and the non-inferiority analysis for clinical outcome

By simulating increased mortality in the BL-group, RADAR remained statistically significant above the 55% threshold up to an increased relative risk of death of 1.5 in the analysis with five clinical outcome categories and up to an increased relative risk of death of 1.25 in the analysis with eight clinical outcome categories (Figure 2). Thus, in spite of this marked increase in mortality in the BL group, RADAR continued to show a favorable outcome for BL. In contrast, the conventional non-inferiority approach already failed to demonstrate non-inferiority after the first step increase of the relative risk of death at a level of 1.125.

**Figure 2.**
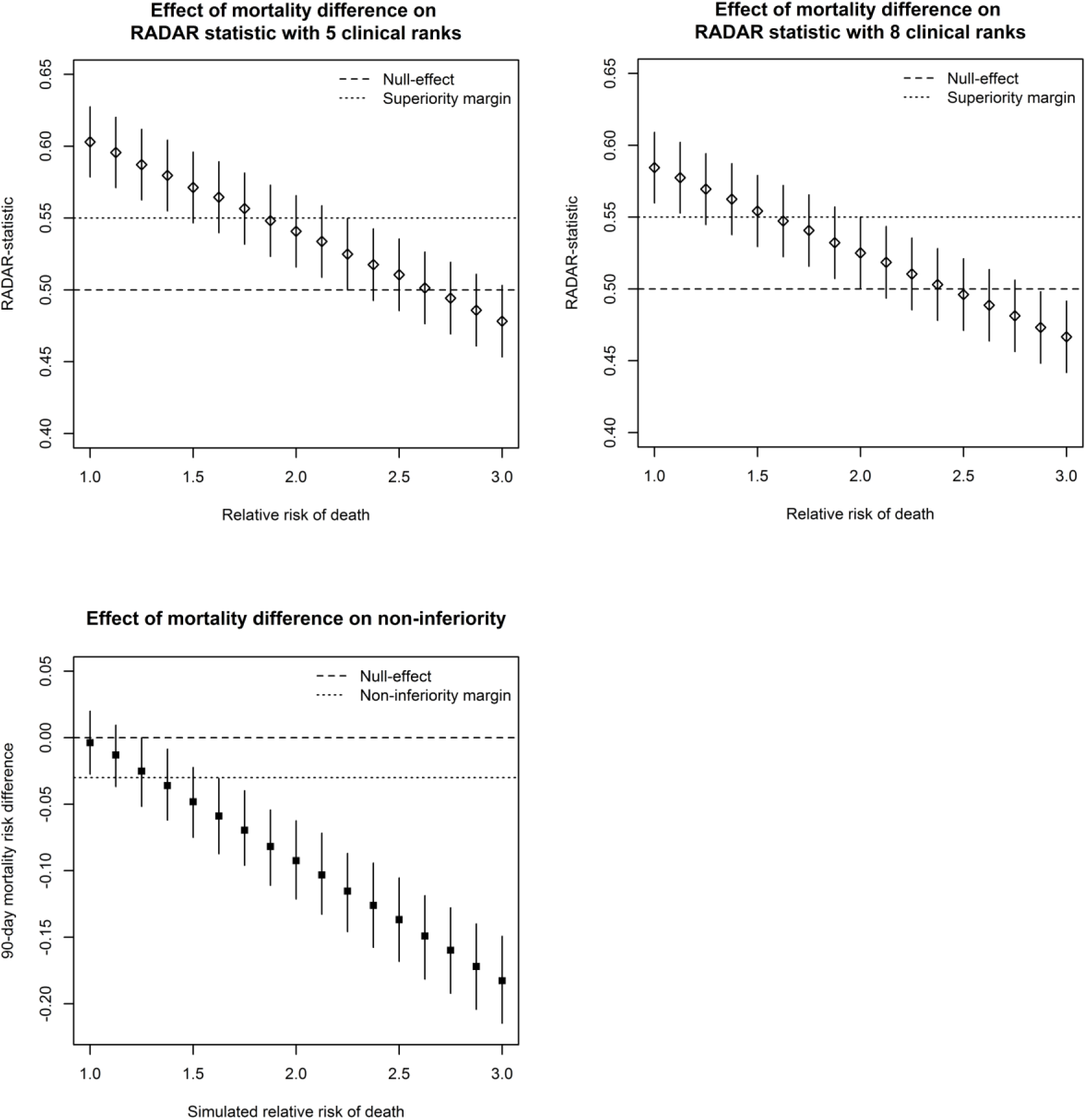
Relation between a simulated increase in mortality and the RADAR-statistic or non-inferiority. The definition of the five and eight clinical outcome categories are defined in table 1.

## DISCUSSION

Application of the recently proposed RADAR on data from a multicenter cluster randomized trial, designed to demonstrate non-inferiority of a strategy of BL for 90-day all-cause mortality in patients with moderate-severe CAP, revealed a significantly more favorable ranking in the BL group compared to the FQL group. In our study, the value of RADAR was shown to be strongly influenced by the number of clinical outcome categories used but not to the number of categories for antibiotic use. Despite the favorable RADAR outcome, we showed that the obtained value of RADAR varies with the number of clinical outcome categories chosen in the design phase. The performed sample size calculations show that RADAR may require fewer than the conventional non-inferiority analysis. However, RADAR and the reduced sample size showed to be insensitive to detecting worse clinical outcomes.

The positive RADAR confirm the absence of differences in clinical outcomes with substantially reduced atypical coverage when using the BL strategy^14^. However, we consider interpretation of RADAR as difficult. With no differences in clinical outcome, the statistic was substantially higher using two clinical categories as compared to five or eight clinical categories. Furthermore, a RADAR-statistic >50% should indicate that the clinical outcome is better and/or that antibiotic use has decreased. However, the simulations with increasing mortality rates in the BL strategy revealed that positive RADAR-statistics can still be found in the setting of a worse clinical outcome. In addition, RADAR represents the probability of more favorable ranking, without quantification of intervention effectiveness.

Our findings demonstrate that the construction of the clinical outcome categories with the hierarchical levels needs to be considered carefully, as it influences both the sample size calculation and the number of predefined clinical outcome categories directly determined the value of RADAR. The latter can be explained by a reduced contribution of antibiotic use to RADAR when more hierarchical levels are used. In the extreme case, if the resolution of clinical outcome is infinite, every patient would get its own clinical outcome category, negating the effect of antibiotic use completely, as this only affects patients that are at the same clinical outcome category. As a result, the value of RADAR can be influenced by choosing the amount of clinical outcome categories, which makes comparison of RADAR-statistics between studies only possible if the same ranking categories have been used.

An attractive argument to adopt RADAR would be a required lower sample size^10^. When comparing the calculated required sample size of RADAR to the original sample size required for the classical non-inferiority design of CAP-START, there was a 61% reduction in required sample size when using five clinical outcome categories but only a 11% reduction when using eight clinical outcome categories. However, as RADAR is a composite endpoint, separate analysis of clinical outcomes is still required to ensure safety of the reduction in antibiotic use, and a lower sample size results in less confidence of safety to the intervention for clinical outcome^12^. This was illustrated by the simulated trial analysis with the lower RADAR sample size, that did not allow the conclusion of non-inferiority for 90-day mortality when the RADAR method required a marked lower sample size, as was the case when using five clinical outcome categories. We, therefore, concur with others that RADAR should not be used as the main criterion to determine the required sample size^12,13^.

Strengths of the current analysis include the application of the CAP-START trial, constituting real-life trial data, to evaluate RADAR. The design of the CAP-START trial is very suitable to evaluate the RADAR method, as the randomized antibiotic strategies allowed for deviation. This resembles daily practice of antibiotic stewardship interventions where deviations from a proposed strategy are frequently justified. Also, the sample size calculations were performed using both the estimated and the actual clinical outcomes and antibiotic use as found in the CAP-START study. Other strengths include the completeness of data collection and the sample size allowing comparison between a conventional non-inferiority and RADAR.

A limitation of the current analysis is that we did not adjust for clustering in the sample size calculations and in the calculation of RADAR. Due to the cross-over design, the intra-cluster correlation of 90-day mortality was negligible in this study, therefore, the impact of not adjusting for clustering was expected to be minimal^14^. However, the intra-cluster correlation of antibiotic use could be higher and taking this into account might have resulted in a higher required sample size for the RADAR method.

To conclude, we present the application of the novel RADAR methodology to analyze antibiotic stewardship interventions on a previously conducted non-inferiority trial. RADAR inventively combines clinical outcome with antibiotic use as a composite endpoint on a patient level. However, we believe that it should not replace non-inferiority of patient-relevant clinical endpoints as the primary analysis. Therefore, we recommend that mortality is used for sample size calculations (as this endpoint requires the largest number of patients) and that mortality and antibiotic use are reported as the two primary endpoints in antibiotic stewardship trials.

